# Alphaproteobacteria facilitate *Trichodesmium* community trimethylamine utilization

**DOI:** 10.1101/2021.03.10.434842

**Authors:** Asa E. Conover, Michael Morando, Yiming Zhao, Jacob Semones, David A. Hutchins, Eric A. Webb

## Abstract

In the surface waters of the warm oligotrophic ocean, filaments and aggregated colonies of the nitrogen (N)-fixing cyanobacterium *Trichodesmium* create microscale nutrient-rich oases. These hotspots fuel primary productivity and harbor a diverse consortium of heterotrophs. Interactions with associated microbiota can affect the physiology of *Trichodesmium*, often in ways that have been predicted to support its growth. Recently, it was found that trimethylamine (TMA), a globally-abundant organic N compound, inhibits N_2_ fixation in cultures of *Trichodesmium* without impairing growth rate, suggesting that *Trichodesmium* receives nitrogen from TMA. In this study, ^15^N-TMA DNA stable isotope probing (SIP) of a *Trichodesmium* enrichment was employed to further investigate TMA metabolism and determine if TMA-N is incorporated directly or secondarily via cross-feeding facilitated by microbial associates. Herein we identify two members of the marine *Roseobacter* clade (MRC) of Alphaproteobacteria as the likely metabolizers of TMA and provide genomic evidence that they converted TMA into a more readily available form of N, e.g., NH_4_^+^, which was subsequently used by *Trichodesmium* and the rest of the community. The results implicate microbiome-mediated carbon (C) and N transformations in modulating N_2_ fixation, and thus highlight the involvement of host-associated heterotrophs in global biogeochemical cycling.

## Introduction

The N-fixing cyanobacterium *Trichodesmium* contributes a substantial portion of the new nitrogen and carbon in the warm oligotrophic ocean (Capone *et al*., 1997; Karl *et al*., 1997; Capone *et al*., 2005; Zehr and Capone, 2020). *Trichodesmium* grows in multicell filaments called trichomes, which can aggregate to form macroscopic colonies. Trichomes and colonies of *Trichodesmium* create micro-scale nutrient hotspots, referred to as the phycosphere or trichosphere, that attract and support the growth of many heterotrophic organisms (e.g., Hmelo *et al*., 2012; Frischkorn *et al*., 2017; Klawonn *et al*., 2020; Zehr and Capone, 2020). Evidence suggests that such heterotrophs may confer physiological benefits to *Trichodesmium*, such as through drawdown of oxygen (an inhibitor of N_2_ fixation), production of CO_2_ (required for photosynthesis), detoxification of radical oxygen species, and production of siderophores and alkaline phosphatases for acquisition of Fe and P, respectively (Paerl *et al*., 1989; Webb *et al*., 2007; Hynes *et al*., 2009; Roe *et al*., 2012; Lee *et al*., 2017; Basu *et al*., 2019). Attempts to culture *Trichodesmium* axenically have been challenging, though a single successful effort in the early 2000s by Waterbury found that axenic cultures of *Trichodesmium erythraeum* IMS101 grow at a reduced rate (personal communication), further suggesting that associated microbiota play an important role in *Trichodesmium* physiology.

The tight association between *Trichodesmium* and its heterotrophic consortium has made it historically difficult to discern which metabolic functions specific taxa perform and what chemical compounds mediate these interactions. Recent studies have suggested that methylated amines (MAs) may play important functions in *Trichodesmium* consortium dynamics. Pade *et al*. (2016) found that *Trichodesmium erythraeum* IMS101 produces the quaternary ammonium compound N,N,N-trimethyl homoserine (or homoserine betaine), a member of the methylamine family, as its main compatible solute to maintain osmotic balance. Walworth *et al*. (2018) showed that TMA, a globally-abundant organic N compound, can support growth of *T. erythraeum* IMS101 in non-axenic culture while suppressing N_2_ fixation, suggesting that *Trichodesmium* can use TMA as a N source. The latter study further found that long-term growth with elevated CO_2_ and limiting nutrients led *Trichodesmium* to downregulate transcription of the N_2_-fixing nitrogenase enzyme and upregulate production of a gene predicted to encode trimethylamine monooxygenase (Tmm) — an enzyme that catalyzes the oxidation of TMA to trimethylamine N-oxide (TMAO), the first step in aerobic TMA catabolism (Chen *et al*., 2011; Lidbury *et al*., 2014; Sun *et al*., 2019). But while *T. erythraeum* IMS101 possesses a likely Tmm homologue, it lacks other known genes required for TMA utilization (Walworth *et al*., 2015). It thus remains unclear if *Trichodesmium* is capable of consuming TMA directly, or if TMA is first metabolized by associated microbiota and then transferred to *Trichodesmium* as an alternate form of nitrogen.

*Trichodesmium* colonies have been found to host likely metabolizers of MAs. *Roseibium* sp. TrichSKD4, a member of the marine *Roseobacter* clade (MRC) of Alphaproteobacteria, was isolated from *Trichodesmium* colonies in defined media and subsequently found to possess the full set of genes required for TMAO catabolism (Lidbury, 2015). However, it lacks the *tmm* gene for conversion of TMA to TMAO (Lidbury, 2015), suggesting potential to benefit from *Trichodesmium* derived Tmm activity.

Chen *et al*. (2012) demonstrate that several members of the MRC can use MAs, including TMA, as a sole N source, and that many have the genetic potential to do so. Lidbury *et al*. (2015) show that TMA catabolism in MRC member *Ruegeria pomeroyi* DSS-3 generates cellular energy and remineralizes ammonium, which supports the growth of co-cultured bacteria. The production of ammonium, a known N source for *Trichodesmium* (Mulholland and Capone, 1999), highlights a potential mechanism whereby *Trichodesmium* may receive N through exogenous TMA metabolism. However, the occurrence of such secondary utilization, and more broadly, the potential influence that recycling of N compounds by heterotrophs has on rates of N_2_ fixation, has yet to be resolved.

In this study, we examine whether *Trichodesmium* receives nitrogen from TMA directly (Figure 1, Option A) or secondarily via conversion to an alternate nitrogen species by a co-cultured heterotroph (Figure 1, Option B). ^15^N-TMA DNA SIP was used to investigate TMA metabolism in non-axenic cultures of *Trichodesmium erythraeum* GBRTRLIN201 (hereafter LIN). Our results suggest that TMA catabolism by co-cultured MRC taxa has the ability to provide *Trichodesmium* with a N source, and thereby modulate N_2_ fixation.

**Figure 1.**
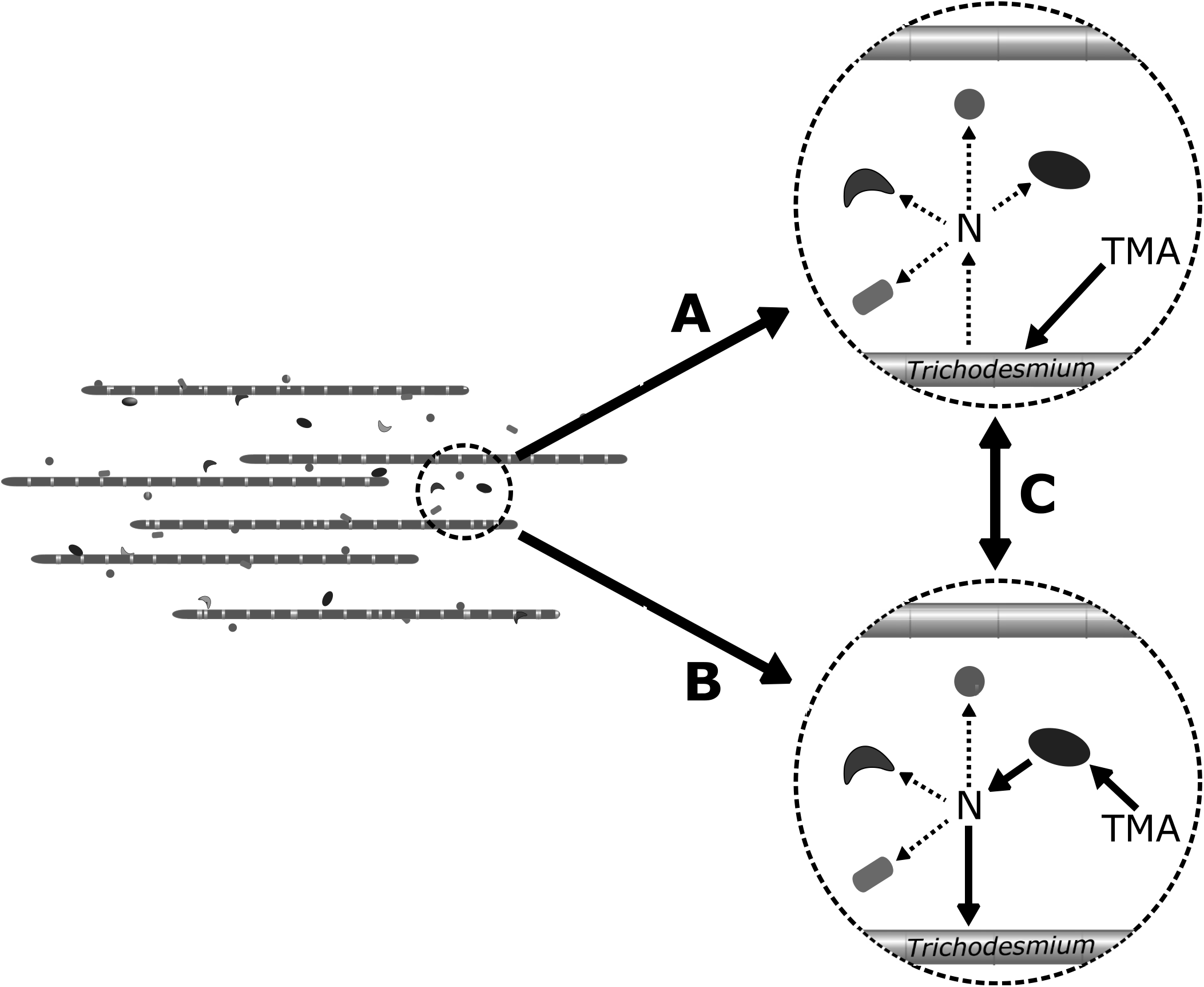
Potential paths of TMA incorporation in the *Trichodesmium* consortium. Option A - *Trichodesmium* metabolizes TMA directly, possibly then releasing a nitrogen compound that is incorporated by associated taxa. Option B -*Trichodesmium* receives TMA-N secondarily following conversion of TMA to an alternate N compound by one or more of the associated taxa. Option C - both *Trichodesmium* and one or more of the associated taxa metabolizes TMA.

## Results

As seen in Walworth *et al*. (2018) with *T. erythraeum* IMS101, the addition of 80 µM TMA suppressed N_2_ fixation by day 1 in both heavy (^15^N) and light (^14^N) treatments of *T. erythraeum* LIN culture, reducing fixation rates to 10 - 30% that of the no-addition control cultures (Figure 2, bottom panel). On day 2, N_2_ fixation remained at about 10% that of the no-addition control in both TMA-addition groups. Fixation rates climbed on day 3 in two of three light TMA-addition cultures but remained suppressed in all others. On day 4, N_2_ fixation increased in all TMA-addition cultures, though still at 50 - 75% that of the no-TMA control cultures. Over the four days, despite the reduced levels of N_2_ fixation, heavy and light TMA-addition cultures grew at similar rates to the no-addition controls, as measured by *in vivo* chlorophyll A fluorescence measurements (Figure 2, top panel).

**Figure 2.**
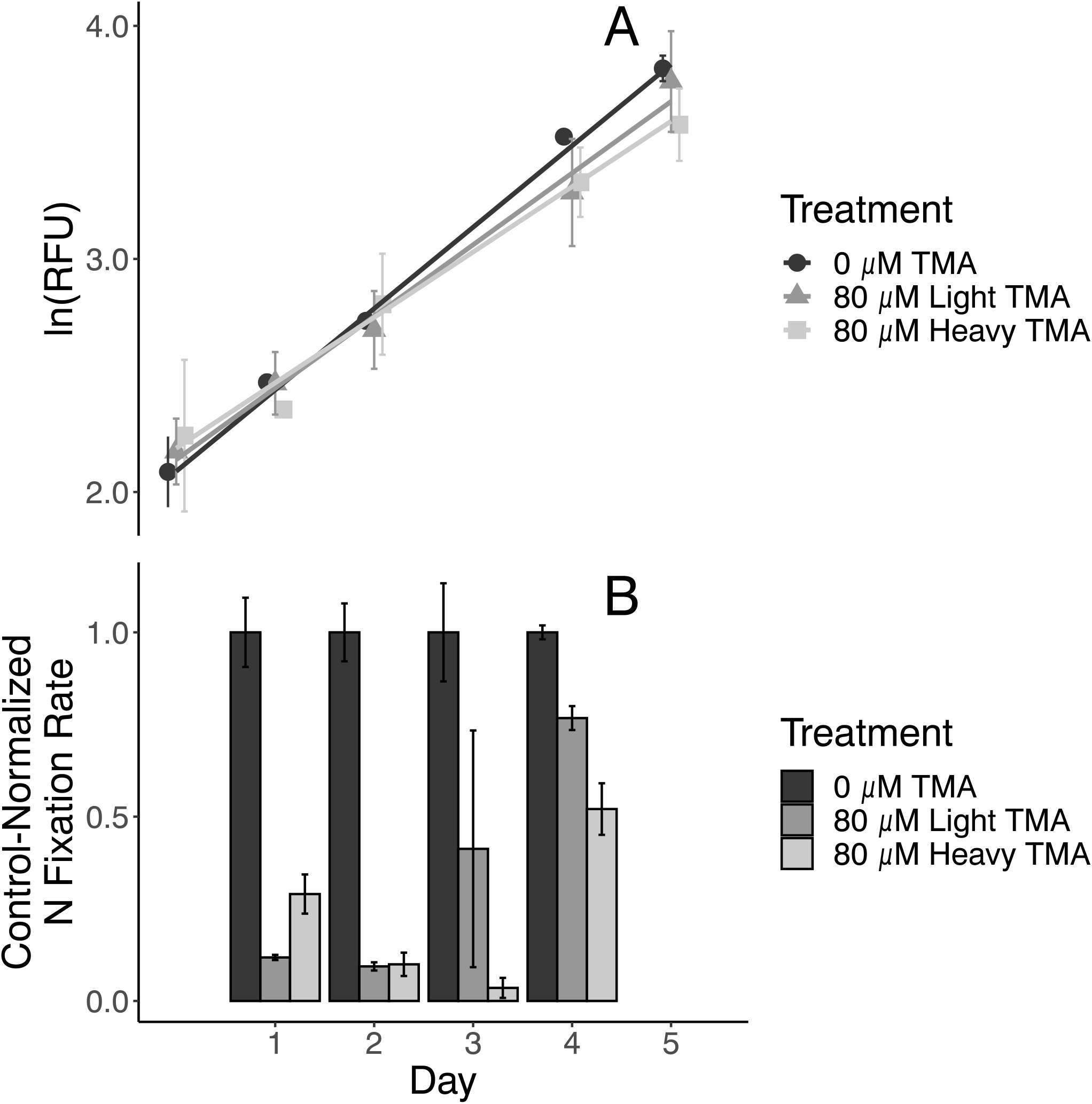
Growth and N_2_ fixation rates for ^14^N TMA, ^15^N TMA, and no TMA control cultures over the four days following TMA addition. A = *Trichodesmium* growth, represented by ln-transformed chlorophyll A fluorescence, measured *in vivo* in relative fluorescence units (RFU). B = rate of N_2_ fixation divided by each culture’s chlorophyll A fluorescence and normalized to the value of the no-addition control treatment. N_2_ fixation rates were measured by gas chromatography using the acetylene reduction proxy method.

^15^N-SIP was used to track utilization of the isotopically-labeled TMA. Density fractionation yielded 18-19 fractions per sample with sufficient DNA for PCR amplification. Metabarcode sequencing of 16S/18S genes generated a total of 1,171,632 reads following denoising and sparsity filtering, with a mean of 15,832 reads per fraction (maximum = 33,064, minimum = 8,280). Twenty bacterial ASVs were common to both treatments and present in all three timepoints. No 18S ASVs passed the filtering threshold. Atom fraction excess was calculated for each ASV using the qSIP function of the HTSSIP R package (Youngblut *et al*., 2018). On day 1, 3 ASVs showed label incorporation—two belonging to Alphaproteobacteria of the *Rhodobacteraceae* family (Seq13, Seq14), and one to a Gammaproteobacterium of unknown family (Seq05). By day 3, all of the community, including *Trichodesmium* (Seq04), showed atom fraction excess values indicative of ^15^N incorporation. On day 4,^15^N enrichment remained high, with atom fraction excess values dropping below the enrichment threshold in just two of the 20 ASVs—Seq18 and Seq19, both belonging to the *Bacillales* order of Firmicutes.

The two day 1 *Rhodobacteraceae* incorporators share 100% identity with V4-V5 region 16S sequences from published genomes available on IMG/M. Specifically, Seq13 is 100% identical to *Mameliella spp*. (10 genomes), while Seq14 is 100% identical to *Phaeobacter gallaeciensis* (7 genomes). Both *Mameliella* and *Phaeobacter* are members of the marine *Roseobacter* clade (MRC) of Alphaproteobacteria.

To determine whether *Mameliella spp*. or *P. gallaeciensis* are indeed present in *Trichodesmium* cultures, genomes of each were obtained from IMG/M and used as read-mapping references for metagenomic reads from cultures of *T. erythraeum* LIN and other *Trichodesmium* strains. Anvi’o visualization of the Bowtie2 read-mapping showed that *Mameliella spp*. genomes were detectable at low concentration in the non-TMA-treated LIN metagenome, but *P. gallaeciensis* genomes were not (Supplemental Figure 2). *Mameliella spp*. was also detected in metagenomes of other strains of *T. erythraeum*, as well as in *T. thiebautii* VI-I (data not shown). It is possible that a relative of *P. gallaeciensis* is present in the enrichments but was not detected in this analysis either because its genome is different from the ones obtained from IMG/M or its abundance was low in the non-TMA-enriched cultures used for sequencing.

To determine whether Seq13 and Seq14 associate with *Trichodesmium* in the field, a BLAST database was constructed using 16S sequences of the *Trichodesmium* consortium collected by Rouco *et al*. (2016) across the Atlantic and Pacific Ocean. Multiple 100% matches to both ASVs were found in colonies of “puff” morphology from the South Pacific, but no identical hits appeared in colonies of other oceanic regions or of the “raft” morphology (Supplemental File 1).

BLASTP was used to assess genomic potential for TMA catabolism in *Mameliella spp*., *T. erythraeum*, and environmental collections of the *Trichodesmium* consortium (Supplemental File 2, Figure 4). All 10 publicly-available *Mameliella* genomes were found to possess likely homologs for the full suite of genes required for the aerobic TMA catabolism pathway described by Lidbury *et al*. (2015) (Figure 4a). In contrast, the three queried strains of *T. erythraeum*, including LIN, were found to possess possible homologs of Tmm and GmaS, but none of the other 12 genes in the pathway. Possible homologs for all 14 genes appeared in at least one of the three metagenomes of *Trichodesmium* colonies collected near Station ALOHA (Figure 4, Supplemental File 2).

## Discussion

Associated microbiota dramatically expand the metabolic potential and functional significance of *Trichodesmium* (Frischkorn *et al*., 2017; Lee *et al*., 2017; Lee *et al*., 2018). As such, the role of the *Trichodesmium* consortium in N cycling stretches beyond N_2_ fixation. High rates of nitrate and nitrite assimilation and reduction are detected in *Trichodesmium* colonies, with negligible detected loss of N to denitrification (Klawonn *et al*., 2020). This network of N transformation processes demonstrates high potential for recycling N, a limiting nutrient for many sympatric, oligotrophic organisms. Metagenomic and transcriptomic surveys of *Trichodesmium* colonies implicate associated heterotrophs in many of these recycling processes (Frischkorn *et al*., 2017; Gradoville *et al*., 2017; Lee *et al*., 2018). Although it has been shown that *Trichodesmium* derives benefit from its microbiota, it is unknown whether it receives N through these recycling processes.

TMA is a useful compound for investigating this question. While cultures of *T. erythraeum* appear to use TMA as a N source in lieu of N_2_ fixation (Figure 2, Walworth *et al*., 2018), the genome of *T. erythraeum* lacks most of the known genes for TMA catabolism (Figure 4, Walworth *et al*., 2015). Therefore, unless *T. erythraeum* metabolizes TMA through a novel biochemical pathway, it must receive TMA-N in an alternate form, i.e., following TMA catabolism by associated microbiota.

In this study, SIP was employed to determine if TMA-N was incorporated directly or secondarily via cross-feeding facilitated by the *Trichodesmium* consortium. Our results revealed that two Alphaproteobacteria of the MRC were ^15^N-enriched before other taxa (Figure 3) and may have converted this N source into a more bioavailable form that was subsequently used by the rest of the community. Additionally, we found genomic evidence that these MRC taxa have the metabolic potential to catabolize TMA, thereby transforming TMA nitrogen to ammonium, further demonstrating the potential for cross-feeding over time (Figure 3 and 4).

**Figure 3.**
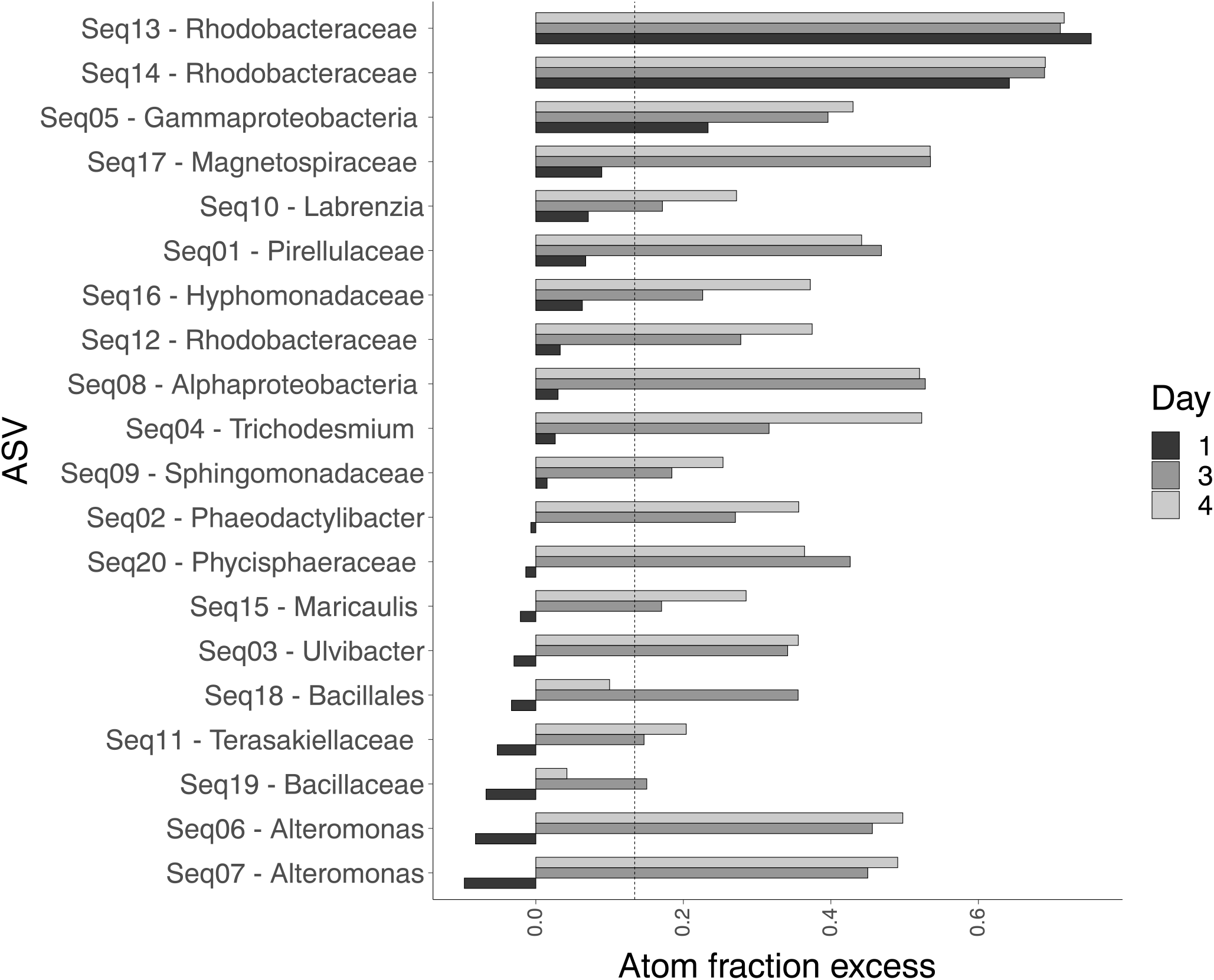
Atom fraction excess of ^15^N for each ASV on days 1, 3, and 4. The dotted vertical line at 0.1340 represents our empirical threshold for incorporation of ^15^N. Each ASV is listed on the X-axis along with the finest-level classification assigned by QIIME2’s q2-feature-classifier plugin at the default confidence threshold of 0.7 using the SILVA132 database.

**Figure 4.**
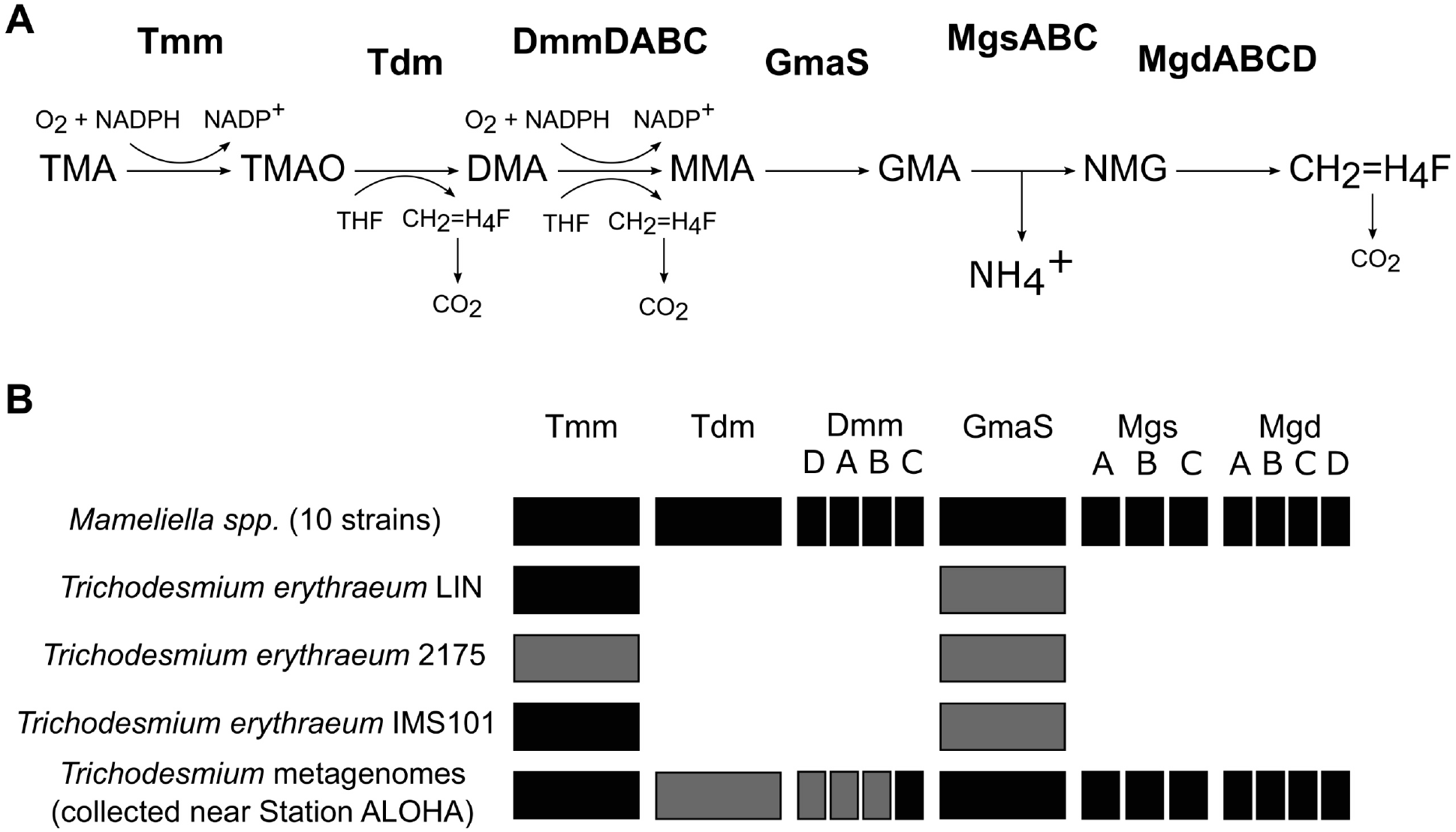
Proposed model for methylated amine catabolism and presence/absence of corresponding proteins in the genomes of *Trichodesmium* and *Mameliella spp*. (a) Proposed MA catabolism pathway as described by Lidbury *et al*. (2015). (b) Presence/absence of gene homologs in *T. erythraeum, Mameliella spp*., and environmental *Trichodesmium* metagenomes based on evidence from BLASTP search for TMA utilization proteins from *R. pomeroyii* DSS-3 (Chen *et al*., 2011; Lidbury *et al*., 2014; Lidbury *et al*., 2015; Lidbury *et al*., 2017). Black rectangles denote BLASTP hits with ≥ 90% coverage and ≥ 50% identity. Gray rectangles denote BLASTP hits with ≥ 90% coverage and ≥ 25% identity or ≥ 50% coverage and ≥ 50% identity. For *Mameliella spp*. (10 strains), rectangles indicate hits in genomes from all 10 strains. For *Trichodesmium* metagenomes (collected near Station ALOHA), rectangles indicate hits in one or more of the three metagenomes. Tmm, TMA monooxygenase (Chen *et al*., 2011); Tdm, TMAO demethylase (Lidbury *et al*., 2014); Dmm, DMA monooxygenase (Lidbury *et al*., 2017); GmaS, gamma-glutamylmethylamide synthetase (Lidbury *et al*., 2015); MgsABC, N-methylglutamate synthase (Lidbury *et al*., 2014); MgdABCD, N-methylglutamate dehydrogenase (Lidbury *et al*., 2014). TMA, trimethylamine; TMAO, trimethylamine N-oxide; DMA, dimethylamine; MMA, monomethylamine; GMA, gamma-glutamylmethylamide; NMG, N-methylglutamate.

The MRC are globally-distributed and abundant in a wide variety of environments—from the coast to the open ocean, and from surface waters to deep seafloor sediments (Buchan *et al*., 2005; Luo and Moran, 2014). Members of the MRC are commonly found in association with phytoplankton (Geng and Belas, 2010) and have been shown to benefit their phytoplankton hosts in some of these relationships, such as through production of growth-stimulating auxins (Seyedsayamdost *et al*., 2011) and protective antibiotics (Rao *et al*., 2007). MRC members typically possess comparatively large bacterial genomes and demonstrate high metabolic versatility (Newton *et al*., 2010). Surveys of the *Trichodesmium* consortium in culture and in the field have regularly found MRC taxa present (Hmelo *et al*., 2012; Rao *et al*., 2015; Rouco *et al*., 2016; Lee *et al*., 2017). Thus, members of the MRC may play an important role in *Trichodesmium* ecology and physiology.

Genes for TMA and TMAO metabolism are common but not universal among the MRC (Chen, 2012; Lidbury *et al*., 2014) and enable some taxa to use TMA as a sole N source (Chen, 2012). Landa *et al*. (2017) suggest that the patchy distribution of TMA utilization genes among the MRC may implicate TMA as a currency of specific relationships between individual roseobacters and phytoplankton species, many of which (including *Trichodesmium*) produce MAs as compatible solutes to maintain osmotic balance in saline environments (Mackay *et al*., 1984; Oren, 1990; Pade *et al*., 2016). These compounds accumulate in high concentrations and represent substantial pools of organic N, which are liberated during cellular lysis (Welsh, 2000).

While the metabolic fate of the major compatible solute in *Trichodesmium*, trimethylated homoserine, remains unknown, other quaternary amines, such as glycine betaine and choline, are known to degrade to TMA (King, 1984; Oren, 1990; Bain *et al*., 2004). Thus, trimethylated homoserine constitutes a likely source of TMA *in situ*, potentially providing an ecological incentive to microbiota that can metabolize TMA and other MAs. Not surprisingly, a member of the MRC isolated from *Trichodesmium* colonies, *Roseibium sp*. TrichSDK4, was found to possess the genes for TMAO metabolism (Lidbury *et al*., 2014).

Lidbury *et al*. (2015) propose a model for TMA catabolism via aerobic oxidation in the MRC member *R. pomeroyi* DSS-3. They demonstrate that this pathway generates ATP and leads to remineralization of ammonium—some of which is kept as an N source while the rest is exported. Our results are consistent with this model: high levels of ^15^N incorporation by the two MRC ASVs on day 1 were followed by incorporation in nearly all of the community, including *Trichodesmium*, on day 3, and further incorporation on day 4. Such widespread incorporation implies conversion of TMA to a more accessible nitrogen species, such as ammonium, as is predicted by the aerobic oxidation pathway (Lidbury *et al*., 2015). This trophic cascade suggests that TMA metabolism in *T. erythraeum* LIN cultures proceeds as outlined in Figure 1, Option B—in which MRC taxa Seq13 and Seq14 perform TMA catabolism and release ammonium, which then serves as an N source for *Trichodesmium* and the rest of the community. In this scenario, the rapid inhibition of N_2_ fixation in *Trichodesmium* apparent on day 1 (Figure 2) is explained by MRC ammonium production.

While the results support this model, they do not rule out the possibility that *Trichodesmium* is also capable of TMA metabolism. It is possible that both MRC and *Trichodesmium* utilize TMA (i.e., Figure 1, Option C) and that we saw MRC incorporation first because they had faster generation times, leading to more rapid incorporation of ^15^N. Indeed, genomic-based estimates collected in the EGGO database predict a maximal doubling time of ∼2.4 hours for *Mameliella alba* CGMCC [GCF_900101505.1] (Weissman *et al*., 2020), while the doubling time of *T. erythraeum* LIN observed here was ∼48 hours. However, if *Trichodesmium* were to utilize TMA directly, it would need to do so by a yet unknown pathway, since *T. erythraeum* LIN (along with IMS101 and all other genome-sequenced strains) lack most of the known genes required for TMA metabolism (Figure 4, Walworth et al., 2018). In contrast, ammonium is a well-documented N source for *Trichodesmium* (Mulholland and Capone, 2000), and as described above, was likely made available via MRC TMA catabolism. Therefore, it appears likely that suppression of N_2_ fixation in cultures of *T. erythraeum* LIN (and by proxy IMS101 as well) following TMA addition was the result of TMA catabolism by co-cultured MRC taxa.

It is not difficult to envision how such an association could prove mutually beneficial. Through TMA catabolism MRC could gain access to the energy and N source offered by methylated amines released by *Trichodesmium*, such as its primary compatible solute, trimethylated homoserine. In turn, *Trichodesmium* could recover remineralized N released by the MRC associates, thereby saving some of the expenses required for N_2_ fixation. Such an association would promote the consortium’s ability to recycle N and conserve resources. The association may also improve the consortium’s ability to acquire new N. Presumably, the MRC routinely benefits by obtaining N released by *Trichodesmium* (both as MAs, but also as ammonium and other forms of dissolved organic nitrogen), as evidenced by their decades-long perseverance in laboratory cultures in N-free media. But when TMA becomes available to the consortium, the N flux goes the other way—from MRC to *Trichodesmium*. Thus, the direction of N transfer within the *Trichodesmium*/MRC association depends on whether the N source is N_2_ or TMA. This flexibility could benefit both partners as it broadens the entire consortium’s nitrogen source niche.

The environmental data assembled here suggests that this relationship may exist beyond the laboratory. The appearance of Seq13 and Seq14 in *Trichodesmium* colonies from the South Pacific sequenced by Rouco *et al*. (2016) indicates that the same or highly similar MRC taxa associate with *Trichodesmium* in natural environments. And though these same MRC taxa do not appear in colonies globally, the presence of likely TMA catabolism genes in metagenomes collected circa Station ALOHA (near Hawaii) suggests that the aerobic TMA oxidation pathway may be functionally conserved within the consortium. Testing whether TMA addition regularly inhibits N_2_ fixation in the field will be important for assessing the implications of these relationships.

The findings presented here are novel, but not without precedent. Frischkorn *et al*. (2018) demonstrate that microbiota can influence *Trichodesmium* N_2_ fixation rates, but the means by which this happens remain uncertain. This study provides the first evidence we know of that microbiome-mediated N recycling can modulate N_2_ fixation rates. Such evidence elevates the significance of the *Trichodesmium*-associated heterotrophic consortium, which likely plays a significant role in global N_2_ fixation patterns.

## Experimental Procedures

### Culture conditions

Cultures of *T. erythraeum* strain GBRTRLIN201 (Bell *et al*., 2005) were grown from a single inoculum and divided into three TMA treatment groups: final concentration 80 µM ‘heavy TMA’ (^15^N), 80 µM ‘light’ TMA (^14^N), and no TMA addition. Triplicates of each treatment were grown in sterile acid-washed 2 L polypropylene baffled Erlenmeyer flasks with 1.6 L of YBC-II artificial seawater medium (Chen *et al*., 1996). Cultures were housed in Percival (I-66 and I-30) incubators at a temperature of 26.0 °C and light intensity of 200 µEinsteins m^-2^s^-1^ with warm and cool white fluorescent lights operating on a 12 hour light : 12 hour dark cycle. They were kept on an orbital shaker operating at 50 rpm to ensure homogeneity of trichomes. Over the next four days, subsamples were withdrawn from each culture daily to monitor N_2_ fixation and growth rate, as well as to collect biomass for DNA extraction.

### Nitrogen fixation

Nitrogenase activity was quantified by the acetylene reduction assay described by Capone and Montoya (2001) as modified by Chappell *et al*. (2010). This assay leverages the nitrogenase enzyme’s ability to reduce triple-bonded acetylene gas (C_2_H_2_) to double-bonded ethylene gas (C_2_H_4_), both of which can be quantified by gas chromatography. In short, a culture sample in a closed container is injected with acetylene gas, after which ethylene production is measured over time and used as a proxy for calculating the sample’s ability to reduce nitrogen gas (N_2_) to ammonium (NH_4_^+^).

In this study, acetylene gas was generated by mixing 15.0 g of calcium carbide with 50 ml of ddH_2_O in a glass sidearm flask. The evolved gas was collected and stored in a rubber bladder. At the midpoint of the incubator’s daylight cycle 20 ml subsamples of each culture were transferred to clear 30 ml polycarbonate centrifuge tubes with silicone-PTFE septa caps (I-Chem, Thermo Fisher Scientific). Caps were tightly screwed and 3 ml of acetylene gas was injected into each. Tubes were inverted gently to mix. 0.5 ml samples of headspace were withdrawn from each sample using a 1.0 ml disposable syringe. Samples were taken hourly over a two-hour period for a total of three time points. Filled syringes were stuck into a rubber stopper prior to analysis to prevent leaking. 0.2 ml from each syringe was injected in a Shimadzu (GC-8a) gas chromatograph equipped with a flame ionization detector and Porapak R column. Peak height readings were integrated using PeakSimple software (SRI Instruments). A 9.0 ppm ethylene standard was used to calibrate peak heights. Ethylene formed was calculated according to Capone and Montoya (2001) with a Bunsen coefficient of 0.084, as calculated according to Breitbarth *et al*. (2004) for samples at 26°C with salinity of ∼35 ppt. Estimates of nitrogen-fixing capacity were calculated assuming an equivalence of 4 moles ethylene to 1 mole nitrogen gas and values were normalized to biomass using relative fluorescence of chlorophyll A (see below).

### Growth data

Culture growth was monitored via relative fluorescence of chlorophyll A, measured *in vivo* using a Trilogy Laboratory Fluorometer (Turner Designs). Following the conclusion of the daily acetylene reduction assay, *in vitro* chlorophyll concentrations were measured for each sample (Welschmeyer, 1994; Wasmund, 2006).

### DNA extraction

Subsamples were collected from each culture daily for DNA extraction — starting with 250 ml on day 1, 200 ml on day 2, 150 ml on day 3, and 100 ml on day 4. The volume was reduced each day to account for the increasing density of biomass in each culture. Cells were collected on 0.2 µm Supor membrane disc filters (Pall Life Sciences) by gentle vacuum filtration. DNA extraction was performed using a DNeasy PowerSoil Pro kit (Qiagen) following the manufacturer’s protocol.

### Density-gradient stable isotope probing

Density-gradient formation, fractionation, and clean-up was performed similarly to Morando and Capone (2018). Briefly, DNA extracts were mixed with a gradient buffer (0.1 M Tris-HCl, 0.1 M KCl, 1 mM EDTA) and cesium chloride (7.163 M) in 3.3 ml polyallomer centrifuge tubes (Beckman Coulter) to a final density of 1.700 g ml^-1^. Tubes were loaded into a TLN-100 near-vertical rotor (Beckman Coulter) and spun for 72 hours at 136,000 × *g*_av_ and 20°C. Tubes were then fractionated in ∼100 µl increments via displacement with mineral oil. Fraction densities were calculated using an AR200 Digital Refractometer (Reichert Technologies). DNA was precipitated from each fraction with PEG-NaCl (30% PEG, 1.6 M NaCl) and linear polyacrylamide (Invitrogen, Thermo Fisher Scientific), washed with 70% ethanol, dried, and resuspended in TE buffer (10 mM Tris-HCl and 1 mM EDTA, pH 8.0). Recovered DNA was quantified using a Qubit fluorometer with a dsDNA BR Assay Kit (Invitrogen, Thermo Fisher Scientific).

### PCR and sequencing

Fractions from Light TMA day 1 and Heavy TMA days 1, 3, and 4 with quantifiable DNA were sent to a commercial vender (Molecular Research LP, MR DNA) for amplification and MiSeq paired-end (2 × 300 bp) sequencing of the 16S V4-V5 hypervariable regions and partial 18S sequences using the 515F-Y / 926R primer pair (515F-Y: 5’-GTGYCAGCMGCCGCGGTAA, 926R: 5’-CCGYCAATTYMTTTRAGTTT) (Parada *et al*., 2016). Library preparation and sequencing was carried out by MR DNA following Illumina library preparation protocols using MiSeq Reagent Kit v3 (Illumina).

Raw 16S/18S sequence data is available in the NCBI Sequence Read Archive with BioProject ID PRJNA703412 and BioSample IDs SAMN18016931:18017004.

### Inferring sequence variants

Exact sequence variants were inferred from raw reads following the Deblur variant of the Fuhrman Lab eASV Pipeline (dx.doi.org/10.17504/protocols.io.vi9e4h6). Primers were removed using cutadapt (Martin, 2011), and reads were sorted into 16S and 18S pools using BBSplit (Bushnell, 2014). Forward and reverse 16S reads were merged using VSEARCH (Rognes *et al*., 2016) and then trimmed to 365 bases. Forward and reverse 18S reads generated with the 515F-Y / 926R primer pair are non-overlapping, and, therefore, could not be merged. Instead, 18S forward reads were trimmed to 220 bases, reverse reads were trimmed to 179 bases, and the two were concatenated. 16S and 18S reads were denoised separately using Deblur (Amir *et al*., 2017) within QIIME 2 (Bolyen *et al*., 2019). Taxonomy was assigned to ASVs by QIIME 2’s q2-feature-classifier plugin (Bokulich *et al*., 2018) using the SILVA 132 database (Quast *et al*., 2013).

### Identifying ^15^N incorporators

Enrichment with ^15^N was determined using the quantitative SIP (qSIP) function of the HTSSIP software package (v1.4.1; Youngblut *et al*., 2018) in R (v3.6.1; R Core Team 2020). qSIP utilizes ASV-specific density curves from labeled and unlabeled treatments to calculate the shift in density in response to isotope labeling for each ASV (Hungate *et al*., 2015). This shift divided by the calculated maximum shift that would occur with 100% enrichment is termed atom fraction excess and was calculated following Hungate *et al*. (2015) with appropriate parameters for ^15^N taken from Morrissey *et al*. (2018). Data were pre-filtered to remove ASVs that constituted less than 0.01% of the total reads for each treatment, since such low read counts preclude the ability to accurately compare density distributions between treatments.

Without sequencing replicates from each day of the experiment, we were unable to calculate confidence intervals for atom fraction excess values. Instead, we assessed the range of potential error in these values by examining the variation in ASV enrichment on day 1 (Figure 3). Since ^15^N incorporation cannot be less than 0, negative atom fraction excess values obtained on day 1 must be generated by variance in the data. To determine a threshold for likely incorporation of ^15^N, we averaged the negative values seen on day 1 (mean = −0.0447), multiplied by 3, and took the absolute value as the sample deviation, similar to Morando and Capone (2016). We concluded that atom fraction excess values above 0.134 g ml^-1^ are greater than what we might obtain from measurement error and are likely generated by true ^15^N enrichment. Regardless of this threshold, it is clear that significant incorporation occurred in the majority of the ASVs analyzed during the course of the experiment (Figure 3).

### Generating *T. erythraeum* LIN metagenome

DNA extraction was carried out as described above. DNA was sent to Novogene (Sacramento, CA) for library preparation using NEBNext DNA Library Prep Kit (300 bp inserts) and Illumina PE150 sequencing. Reads were processed as described in the “Genome Extraction from Shotgun Metagenome Sequence Data” narrative (https://kbase.us/n/33233/351/) constructed at KBase (Arkin *et al*., 2018) as follows: Reads were quality-checked with FastQC (v0.11.5; Andrew, 2010) and linkers were trimmed using Trimmomatic (v0.36; Bolger *et al*., 2014). Contigs were assembled with metaSPAdes (v3.12.0; Nurk *et al*., 2017) and binned with MaxBin2 (v2.2.4; Wu *et al*., 2016) using kmer frequencies and read depth. PROKKA (v1.12; Seemann, 2014). was used to annotate the assembled bins.

Raw metagenomic sequence data is available in the NCBI Sequence Read Archive with BioSample ID SAMN18208363 within BioProject ID PRJNA703412. The *T. erythraeum* LIN metagenome-assembled genome is available in Supplemental File S4.

### Search for *Mameliella spp*. and *P. gallaeciensis* in *T. erythraeum* LIN metagenome

Ten genomes of *Mameliella spp*. and seven of *P. gallaeciensis* with 16S sequences 100% identical to Seq13 and Seq14, respectively, were identified using BLASTN searching of the JGI IMG/M database (Chen *et al*., 2020) (accessed March 05, 2021). Genomes were downloaded and checked for similarity using FastANI (Jain *et al*., 2018), and highly similar genomes (most with >99% average nucleotide identity) were dereplicated. To avoid search redundancy and random read mapping, the two most divergent genomes of each taxon were selected as representative strains—these were *Mameliella alba* F15 (ID 2788500142), *Mameliella sp*. LZ28 (ID 2891139733), *P. gallaeciensis* P128 (ID 2814123083), and *P. gallaeciensis* DSM26640 (ID 2558309061). Bowtie2 (Langmead and Salzberg, 2012) was used to map metagenomic reads from *T. erythraeum* LIN to these representative genomes. Samtools (Li *et al*., 2009) was used to screen out poor scoring reads in the SAM file and prepare the Bowtie2 output for visualization with Anvi’o (v6.2; Eren *et al*., 2015).

### Search for Seq13 and Seq14 among *Trichodesmium* consortium in the field

To determine if TMA utilizing MRC strains (Seq13 and 14 from LIN) co-occur with natural *Trichodesmium* samples, we made a custom BLAST database of all 16S SSU V4 amplicon reads from a global *Trichodesmium* colony microbial diversity study (Rouco *et al*., 2016). Briefly, sequences were obtained in FASTA format from the NCBI Sequence Read Archive using fastq-dump (https://ncbi.github.io/sra-tools/) and then used to generate a custom BLASTN database in Geneious Prime (www.geneious.com). MegaBLASTN default settings were used and only hits greater than 99.6% across the whole V4 region were reported.

### Surveying genomic TMA catabolism potential

We searched for TMA catabolism proteins in the genomes of *Mameliella spp*. (there are 10 publicly available genomes on IMG/M) and *T. erythraeum* (strains LIN, IMS101, and 2175), as well as in environmental metagenomes generated from *Trichodesmium* colonies (using 3 published from samples collected near Station ALOHA). BLASTP was used to search each genome for the 14 TMA catabolism proteins using reference sequences from *R. pomeroyi* DSS-3, in which most of the proteins were identified (Chen, 2012; Lidbury *et al*., 2014; Lidbury *et al*., 2017). With the exception of *T. erythraeum* LIN, the genome for which was generated in this study, BLASTP searches were conducted using JGI’s IMG/M database. To query the genome of *T. erythraeum* LIN, a custom BLAST+ 2.10.0 database was constructed (Camacho *et al*., 2009).

## Acknowledgements

We would like to thank Doug Capone, John Heidelberg, Jed Fuhrman, and the members of their labs for sharing equipment and space with us. Additionally, we thank Jesse McNichol for guidance with sequence processing and to Mike Lee and Maggi Mars Brisbin for invaluable guidance and conversations. This research was supported by funding from the National Science Foundation (For DAH and EAW: OCE 1657757 and OCE 1851222).

