## Supplementary figures and images for "Alphaproteobacteria facilitate *Trichodesmium* community trimethylamine utilization"

### Supplemental Fig. S1

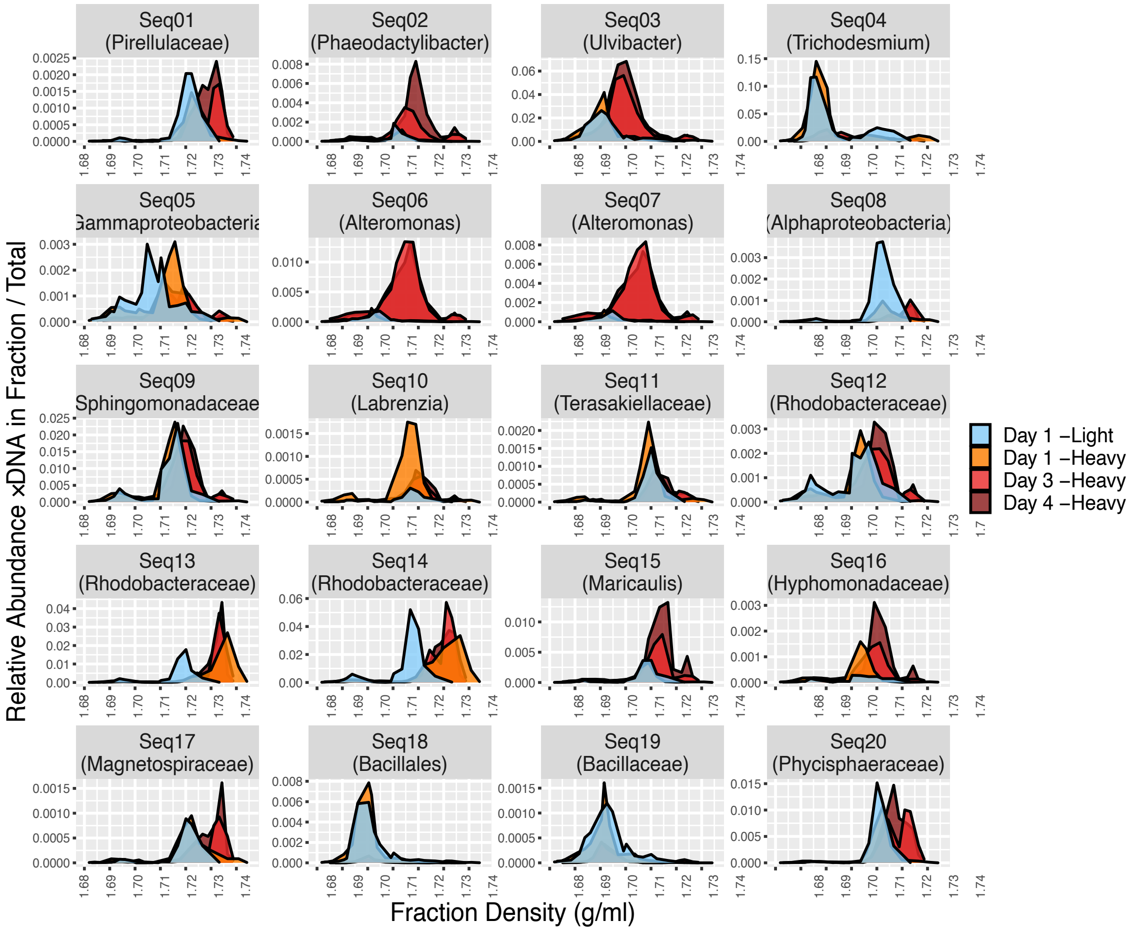

### Supplemental Fig. S2

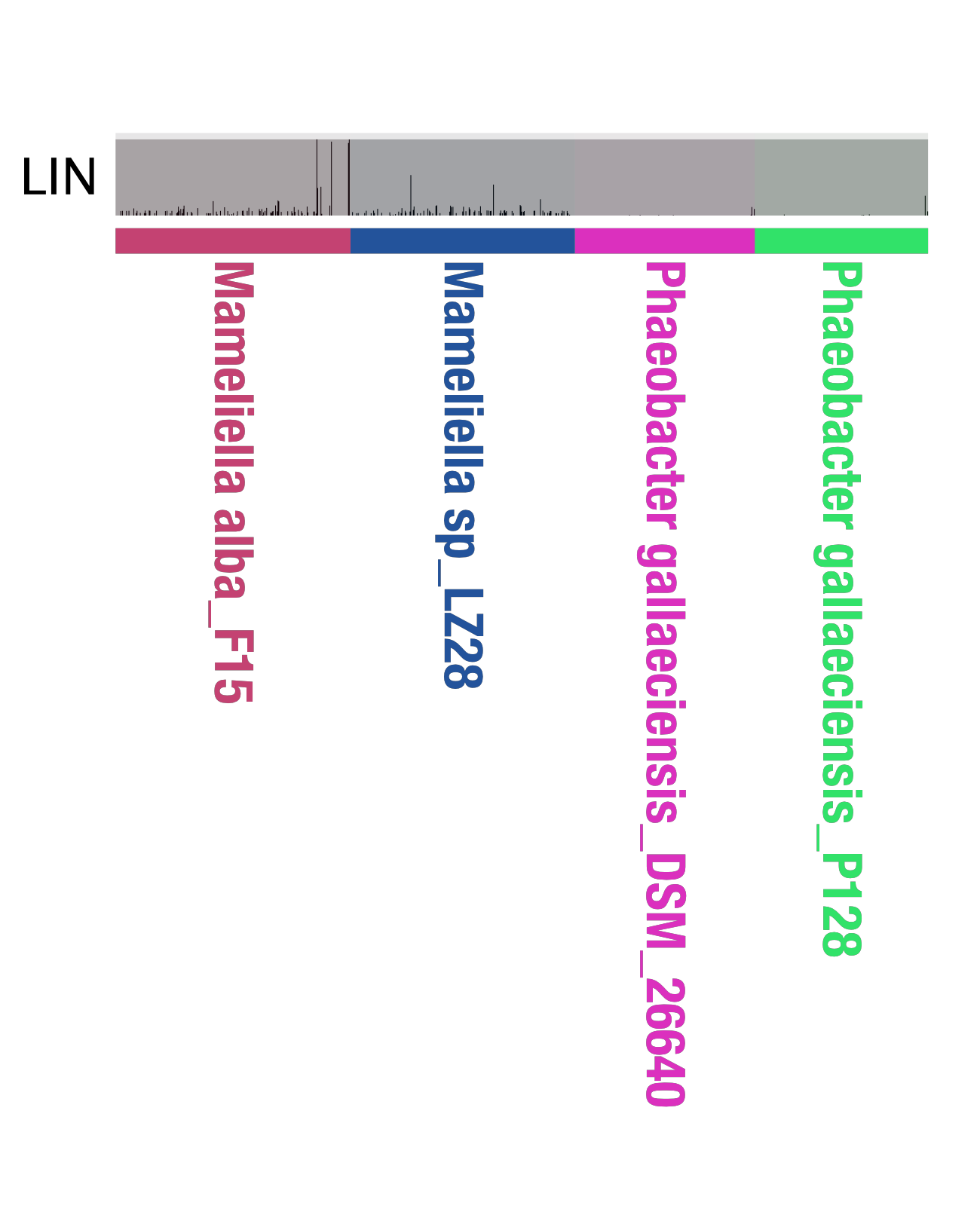
