## Supplemental File S3 for "Alphaproteobacteria facilitate *Trichodesmium* community trimethylamine utilization": qSIP_atom_excess_sourcecode.R

```
#' Calculate G+C from unlabeled buoyant density
#'
#' See Hungate et al., 2015 for more details
#'
#' @param Wlight  A vector with >=1 weighted mean BD from
#' 'light' gradient fractions
#' @return numeric value (fractional G+C; Gi)
#'
calc_Gi = function(Wlight){
  if(length(Wlight) > 1){
    Gi = sapply(Wlight, calc_Gi)
  } else{
    Gi = 1 / 0.083506 * (Wlight - 1.646057)
  }
  return(Gi)
}

#' Calculate the theoretical maximum molecular weight of fully-labeled DNA
#'
#' See Hungate et al., 2015 for more details
#'
#' @param Mlight  The molecular wight of unlabeled DNA
#' @param isotope  The isotope for which the DNA is labeled with ('13C' or '18O')
#' @param Gi  The G+C content of unlabeled DNA
#' @return numeric value: maximum molecular weight of fully-labeled DNA
#'
#'

calc_Mheavymax = function(Mlight, isotope='13C', Gi=NA){
  isotope = toupper(isotope)
  if(isotope=='13C'){
    Mhm = -0.4987282 * Gi + 9.974564 + Mlight
  } else
    if(isotope=='18O'){
      Mhm = 12.07747 + Mlight
    } else 
      if(isotope=='15N'){
        Mhm = 0.5024851 * Gi + 3.517396 + Mlight
        } else {
          stop('isotope not recognized')
    }
  return(Mhm)
}

#' Calculate atom fraction excess
#'
#' See Hungate et al., 2015 for more details
#'
#' @param Mlab  The molecular wight of labeled DNA
#' @param Mlight  The molecular wight of unlabeled DNA
#' @param Mheavymax  The theoretical maximum molecular weight of fully-labeled DNA
#' @param isotope  The isotope for which the DNA is labeled with ('13C' or '18O')
#' @return numeric value: atom fraction excess (A)
#'
calc_atom_excess = function(Mlab, Mlight, Mheavymax, isotope='13C'){
  isotope=toupper(isotope)
  if(isotope=='13C'){
    x = 0.01111233
  } else
    if(isotope=='18O'){
      x = 0.002000429
    } else 
      if(isotope=='15N'){
        x = 0.003663004
      } else {
        stop('isotope not recognized')
      }
  A = (Mlab - Mlight) / (Mheavymax - Mlight) * (1 - x)
  return(A)
}

#' Reformat a phyloseq object of qSIP_atom_excess analysis
#'
#' @param physeq  A phyloseq object
#' @param control_expr  An expression for identifying unlabeled control
#' samples in the phyloseq object (eg., "Substrate=='12C-Con'")
#' @param  treatment_rep  Which column in the phyloseq sample data designates
#' replicate treatments
#' @return numeric value: atom fraction excess (A)
#'
qSIP_atom_excess_format = function(physeq, control_expr, treatment_rep){
  # formatting input
  cols = c('IS_CONTROL', 'Buoyant_density', treatment_rep)
  df_OTU = phyloseq2table(physeq,
                          include_sample_data=TRUE,
                          sample_col_keep=cols,
                          control_expr=control_expr)
  
  # removing 'infinite' BD values
  tmp = colnames(df_OTU)
  df_OTU = df_OTU %>%
    dplyr::mutate_(Buoyant_density = "as.numeric(as.character(Buoyant_density))",
                   Count = "as.numeric(as.character(Count))") %>%
    dplyr::filter_('! is.infinite(Buoyant_density)') %>%
    dplyr::filter_('! is.na(Buoyant_density)') %>%
    as.data.frame
  colnames(df_OTU) = tmp
  
  # return
  return(df_OTU)
}


#' Calculate atom fraction excess using q-SIP method
#'
#' @param physeq  A phyloseq object
#' @param control_expr  Expression used to identify control samples based on sample_data.
#' @param treatment_rep  Which column in the phyloseq sample data designates
#' replicate treatments
#' @param isotope  The isotope for which the DNA is labeled with ('13C' or '18O')
#' @param df_OTU_W  Keep NULL
#'
#' @return A list of 2 data.frame objects. 'W' contains
#' the weighted mean buoyant density (W) values for each OTU in
#' each treatment/control. 'A' contains the atom fraction excess
#' values for each OTU. For the 'A' table, the 'Z' column is buoyant
#' density shift, and the 'A' column is atom fraction excess.
#'
#' @export
#'
#' @examples
#' # tranforming values
#' physeq_rep3_t = OTU_qPCR_trans(physeq_rep3, physeq_rep3_qPCR)
#'
#' \dontrun{
#' # BD shift (Z) & atom excess (A)
#' atomX = qSIP_atom_excess(physeq_rep3_t,
#'                          control_expr='Treatment=="12C-Control"',
#'                          treatment_rep='Replicate')
#' }
#'
qSIP_atom_excess = function(physeq,
                            control_expr,
                            treatment_rep=NULL,
                            isotope='13C',
                            df_OTU_W=NULL){
  # formatting input
  if(is.null(df_OTU_W)){
    no_boot = TRUE
  } else {
    no_boot = FALSE
  }
  
  if(no_boot){
    df_OTU = qSIP_atom_excess_format(physeq, control_expr, treatment_rep)
    if(nrow(df_OTU) == 0){
      stop('No rows in OTU table after qSIP_atom_excess_format(). Check control_exp & treatment_rep')
    }
    
    # BD shift (Z)
    df_OTU_W = df_OTU %>%
      # weighted mean buoyant density (W)
      dplyr::mutate_(Buoyant_density = "as.numeric(as.character(Buoyant_density))",
                     Count = "as.numeric(as.character(Count))") %>%
      dplyr::group_by_('IS_CONTROL', 'OTU', treatment_rep) %>%
      dplyr::summarize_(W = "stats::weighted.mean(Buoyant_density, Count, na.rm=TRUE)") %>%
      dplyr::ungroup()
  }
  
  df_OTU_s = df_OTU_W %>%
    # mean W of replicate gradients
    dplyr::group_by_('IS_CONTROL', 'OTU') %>%
    dplyr::summarize_(Wm = "mean(W, na.rm=TRUE)") %>%
    # BD shift (Z)
    dplyr::group_by_('OTU') %>%
    dplyr::mutate_(IS_CONTROL = "ifelse(IS_CONTROL==TRUE, 'Wlight', 'Wlab')") %>%
    tidyr::spread("IS_CONTROL", "Wm") %>%
    dplyr::mutate_(Z = "Wlab - Wlight") %>%
    dplyr::ungroup()
  
  # atom excess (A)
  ## pt1
  dots = list(~calc_Gi(Wlight))
  dots = stats::setNames(dots, "Gi")
  df_OTU_s = df_OTU_s %>%
    mutate_(.dots=dots) %>%
    mutate_(Mlight = "0.496 * Gi + 307.691")
  ## pt2
  MoreArgs = list(isotope=isotope)
  dots = list(~mapply(calc_Mheavymax, Mlight=Mlight, Gi=Gi, MoreArgs=MoreArgs))
  dots = stats::setNames(dots, "Mheavymax")
  df_OTU_s = df_OTU_s %>%
    dplyr::mutate_(.dots=dots)
  ## pt3
  dots = list(~mapply(calc_atom_excess, Mlab=Mlab, Mlight=Mlight,
                      Mheavymax=Mheavymax, MoreArgs=MoreArgs))
  dots = stats::setNames(dots, "A")
  df_OTU_s = df_OTU_s %>%
    dplyr::mutate_(Mlab = "(Z / Wlight + 1) * Mlight") %>%
    dplyr::mutate_(.dots=dots)
  
  ## flow control: bootstrap
  if(no_boot){
    return(list(W=df_OTU_W, A=df_OTU_s))
  } else {
    return(df_OTU_s)
  }
}


# sampling with replacement from control & treatment for each OTU
sample_W = function(df, n_sample){
  n_light = n_sample[1]
  n_lab = n_sample[2]
  # parsing df
  df_light = df[df$IS_CONTROL==TRUE,]
  df_lab = df[df$IS_CONTROL==FALSE,]
  # sampling
  if(length(df_light$W) > 1){
    W_light = base::sample(df_light$W, n_light, replace=TRUE)
  } else {
    W_light = rep(df_light$W, n_light)
  }
  if(length(df_lab$W) > 1){
    W_lab = base::sample(df_lab$W, n_lab, replace=TRUE)
  } else {
    W_lab = rep(df_lab$W, n_lab)
  }
  # creating new data.frames
  df_light = data.frame(IS_CONTROL=TRUE,
                        #OTU=rep(otu, n_light),
                        W = W_light)
  df_lab = data.frame(IS_CONTROL=FALSE,
                      #OTU=rep(otu, n_lab),
                      W = W_lab)
  return(rbind(df_light, df_lab))
}


# shuffling weighted mean densities (W)
.qSIP_bootstrap = function(atomX,
                           isotope='13C',
                           n_sample=c(3,3),
                           bootstrap_id = 1){
  # making a new (subsampled with replacement) dataset
  n_sample = c(3,3)  # control, treatment
  dots = stats::setNames(list(~lapply(data, sample_W, n_sample=n_sample)), "ret")
  df_OTU_W = atomX$W %>%
    dplyr::group_by_("OTU") %>%
    tidyr::nest() %>%
    dplyr::mutate_(.dots=dots) %>%
    dplyr::select_("-data") %>%
    tidyr::unnest()
  
  # calculating atom excess
  atomX = qSIP_atom_excess(physeq=NULL,
                           df_OTU_W=df_OTU_W,
                           control_expr=NULL,
                           treatment_rep=NULL,
                           isotope=isotope)
  atomX$bootstrap_id = bootstrap_id
  return(atomX)
}


#' Calculate bootstrap CI for atom fraction excess using q-SIP method
#'
#' @param atomX  A list object created by \code{qSIP_atom_excess()}
#' @param isotope  The isotope for which the DNA is labeled with ('13C' or '18O')
#' @param n_sample  A vector of length 2.
#' The sample size for data resampling (with replacement) for 1) control samples
#' and 2) treatment samples.
#' @param n_boot  Number of bootstrap replicates.
#' @param a  A numeric value. The alpha for calculating confidence intervals.
#' @param parallel  Parallel processing. See \code{.parallel} option in
#' \code{dplyr::mdply()} for more details.
#'
#' @return A data.frame of atom fraction excess values (A) and
#' atom fraction excess confidence intervals.
#'
#' @export
#'
#' @examples
#' # tranforming values
#' physeq_rep3_t = OTU_qPCR_trans(physeq_rep3, physeq_rep3_qPCR)
#'
#' \dontrun{
#' # BD shift (Z) & atom excess (A)
#' atomX = qSIP_atom_excess(physeq_rep3_t,
#'                         control_expr='Treatment=="12C-Con"',
#'                         treatment_rep='Replicate')
#'
#' # bootstrapping in parallel
#' doParallel::registerDoParallel(2)
#' df_atomX_boot = qSIP_bootstrap(atomX, parallel=TRUE)
#' head(df_atomX_boot)
#' }
#'
qSIP_bootstrap = function(atomX, isotope='13C', n_sample=c(3,3),
                          n_boot=10, parallel=FALSE, a=0.1){
  # atom excess for each bootstrap replicate
  df_boot_id = data.frame(bootstrap_id = 1:n_boot)
  df_boot = plyr::mdply(df_boot_id, .qSIP_bootstrap,
                        atomX = atomX,
                        isotope=isotope,
                        n_sample=n_sample,
                        .parallel=parallel)
  
  # calculating atomX CIs for each OTU
  mutate_call1 = lazyeval::interp(~ stats::quantile(A, a/2, na.rm=TRUE),
                                  A = as.name("A"))
  mutate_call2 = lazyeval::interp(~ stats::quantile(A, 1-a/2, na.rm=TRUE),
                                  A = as.name("A"))
  dots = stats::setNames(list(mutate_call1, mutate_call2), c("A_CI_low", "A_CI_high"))
  df_boot = df_boot %>%
    dplyr::group_by_("OTU") %>%
    dplyr::summarize_(.dots=dots)
  
  # combining with atomX summary data
  df_boot = dplyr::inner_join(atomX$A, df_boot, c('OTU'='OTU'))
  return(df_boot)
}
```
