## Supplemental File S3 for "Alphaproteobacteria facilitate *Trichodesmium* community trimethylamine utilization": TMA_SIP_allscripts.html

##### Asa Conover

#### 3/7/2021

### Setting up workspace

```
knitr::opts_chunk$set(cache = TRUE, results='hold')
library(phyloseq)
library(reshape2)
library(tidyverse)
library(dplyr)
library(HTSSIP)
library(doParallel)
library(DESeq2)
library(ggplot2)
library(grid)
library(gridExtra)
```

### Physiological data: N fixation + growth rate

```
#read in control-normalized N fixation data
nfix <- read.table("nfix_melt_norm.txt", header=TRUE, check.names = F, na.strings="", sep="\t")
nfix$Treatment <- factor(nfix$Treatment, levels=unique(nfix$Treatment))

#read in ln-transformed RFU data
growth <- read.table("ln_RFU_growth.txt", header=TRUE, check.names = F, na.strings="", sep="\t")
growth$Treatment <- factor(growth$Treatment, levels=unique(growth$Treatment))

#create N fixation plot
nfix_plot <- ggplot(nfix, aes(x=Day, y=nfix, fill=Treatment)) + 
  geom_bar(stat="identity", color="black", 
           position=position_dodge(0.9)) +
  geom_errorbar(aes(ymin=nfix-sd, ymax=nfix+sd), width=.2,
                 position=position_dodge(.9)) +
  labs(y="Control-Normalized \nN Fixation Rate") +
  theme_classic() +
  theme(text = element_text(size=20)) +
  scale_x_continuous(name = "Day", limits = c(-0.1,5.5), breaks = seq(1, 6, by = 1)) +
  scale_y_continuous(breaks = seq(0, 1, by = 0.5)) +
  scale_fill_grey() +
  annotate("text", x = 4.5, y = 1.1, label= "B", size = 10)

#create growth curve
growth_plot <- ggplot(growth, aes(x=Day, y=RFU, color=Treatment, shape=Treatment)) + 
  geom_point(stat="identity", size=4, position=position_dodge(0.25)) +
  geom_errorbar(aes(ymin=RFU-sd, ymax=RFU+sd), width=.2, position=position_dodge(0.25)) +
  geom_smooth(method='lm', se = FALSE) +
  labs(y="\nln(RFU)") +
  theme_classic() +
  theme(text = element_text(size=20)) +
  theme(axis.line.x = element_blank(), axis.ticks.x = element_blank(), axis.title.x = element_blank(), axis.text.x = element_blank()) +
  scale_x_continuous(name = "Day", limits = c(-0.1,5.5), breaks = seq(1, 4, by = 1)) +
  scale_y_continuous(labels = scales::number_format(accuracy = 0.1), breaks = seq(2, 5, by = 1)) +
  scale_color_grey() +
  annotate("text", x = 4.5, y = 4, label= "A", size = 10)

#put the two plots together to share x-axis
grid.newpage()
gridplot <- grid.arrange(growth_plot, nfix_plot)
```

```
## `geom_smooth()` using formula 'y ~ x'
```

```
#save the figure
#ggsave(filename="Figure2_growth-rate_n-fixation.eps", plot= gridplot, device = "eps", dpi= 400, width = 8, height = 8 )
```

### SIP data:

##### Preparing sequencing data, plotting ASV density distributions, using HTSSIP to perform qSIP calculations

#### Importing sequence data

```
#read in 16S and 18S count tables and taxonomies
ps_16S <- import_biom("all-16S-seqs.with-tax.biom")
ps_18S <- import_biom("all-18S-seqs.with-PR2-tax.biom")

#read in metadata
sample_metadata <- import_qiime_sample_data("sample-metadata_HTSSIP.tsv")
sample_metadata$Buoyant_density <- as.numeric(as.character(sample_metadata$Buoyant_density)) #making the Buoyant_density column type 'numeric' rather than a factor

#merging euk and prok data
##first merging count tables
prok_tab <- data.frame(otu_table(ps_16S),check.names=F) 
euk_tab <- data.frame(otu_table(ps_18S), check.names=F)
all_tab <- bind_rows(prok_tab, euk_tab)
all_tab[is.na(all_tab)] <- 0

##next merging taxonomy data
all_taxa <- bind_rows(data.frame(tax_table(ps_16S)), data.frame(tax_table(ps_18S)))

##creating phyloseq object
ps_all <- phyloseq(otu_table(all_tab, taxa_are_rows = TRUE),tax_table(as.matrix(all_taxa)), sample_data(sample_metadata))

#removing in silico mock 16S communities
ps_all <- subset_samples(ps_all, !(sample_names(ps_all) %in% c("16s-mock-staggered-insilico-v2", "16s-mock-even-insilico-v2")) )

# relative abundance transformation
ps_RA <- transform_sample_counts(ps_all, function(x) x/sum(x))
```

#### Prevalence/Abundance filtering

```
#creating quick reference to metadata table 
m = sample_data(ps_RA)

#gather a list of the ASVs that constitute at least 0.1% of the total reads in all timepoints
ps_D1_RA = prune_samples((m$Substrate=='Light' & m$Day==1) | (m$Substrate=='Heavy' & m$Day==1), ps_RA)
D1_sums <- sort(taxa_sums(ps_D1_RA), decreasing = TRUE)
D1_filt <- D1_sums[D1_sums > 0.001*(sum(D1_sums))]

ps_D3_RA = prune_samples((m$Substrate=='Light' & m$Day==1) | (m$Substrate=='Heavy' & m$Day==3), ps_RA)
D3_sums <- sort(taxa_sums(ps_D3_RA), decreasing = TRUE)
D3_filt <- D3_sums[D3_sums > 0.001*(sum(D3_sums))]

ps_D4_RA = prune_samples((m$Substrate=='Light' & m$Day==1) | (m$Substrate=='Heavy' & m$Day==4), ps_RA)
D4_sums <- sort(taxa_sums(ps_D4_RA), decreasing = TRUE)
D4_filt <- D4_sums[D4_sums > 0.001*(sum(D4_sums))]

keepers <- Reduce(intersect, list(names(D1_filt),names(D3_filt),names(D4_filt)))

#manually removing a couple ASVs that are likely contaminants — both have low read counts and implausible density distributions, sporadically appearing across fractions
keepers2 <- keepers[!keepers %in% c("7707aed3364abcbcf1e0c14379673e0c","25aaa1a61e3d99eda26f5b307cf5c39a")]

#creating filtered phyloseq object
ps_filt_all <- prune_taxa(keepers2, ps_all)
```

#### Giving ASVs meaningful names

```
#read in renaming key
renaming_key <- read.table("ASV_naming_key.txt", header=TRUE, row.names=1,check.names = F, na.strings="", sep="\t")
taxa_names(ps_filt_all) <- renaming_key[taxa_names(ps_filt_all),]
```

#### Plotting ASV density distributions

```
#transforming phyloseq object counts to relative abundance 
ps_filt_RA <- transform_sample_counts(ps_filt_all, function(x) x/sum(x))

#creating a table formatted for ggplot2 with relative abundance along with fraction sample name
table_for_plot <- data.frame(otu_table(ps_filt_RA), check.names = FALSE)
table_for_plot$seq <- row.names(table_for_plot)
table_for_plot.g <- gather(table_for_plot, Sample, Proportion, -seq)

#splitting fraction sample name to get number only
fraction_num <- strsplit(as.character(table_for_plot.g$Sample), ".", fixed = TRUE)
fraction_num2 <- sapply(fraction_num, function(x) x[2])
table_for_plot.g$Fraction <- as.numeric(fraction_num2)   

#adding in the metadata to get density values
table_for_plot.g2 <- merge(table_for_plot.g, data.frame(sample_data(ps_filt_RA), check.names = FALSE), by.x = "Sample", by.y = "X.SampleID")

#creating "ProportionScaled" --> multiplying ASV relative abudance by proportion of DNA in the fraction compared to the whole
table_for_plot.g2$ProportionScaled <- table_for_plot.g2$Proportion * as.numeric(as.character(table_for_plot.g2$Fraction.DNA.proportion))

#formatting text for legend
table_for_plot.g2$name <- paste("Day", table_for_plot.g2$Day, "-", table_for_plot.g2$Substrate)
table_for_plot.g2$name <- factor(table_for_plot.g2$name, levels=rev(unique(table_for_plot.g2$name)))

table_for_plot.g2$Buoyant_density <- as.numeric(table_for_plot.g2$Buoyant_density)

ggplot(table_for_plot.g2, aes(x=Buoyant_density, y=ProportionScaled, fill=name)) +
  geom_area(aes(fill=name), color = "black", position = "identity", alpha=0.8) +
  scale_x_continuous(limits = c(1.680,1.7455), breaks = seq(1.680,1.7455,0.010)) +
  theme(text = element_text(size=20), axis.text = element_text(size=10), axis.text.x=element_text(angle=90, vjust=0.4, hjust=1), legend.title=element_blank()) +
  labs(x="Fraction Density (g/ml)", y="Relative Abundance × DNA in Fraction / Total") +
  guides(fill = guide_legend(reverse = TRUE)) +
  scale_fill_manual(values = c("firebrick4","firebrick2","darkorange1","lightskyblue")) +
  facet_wrap( ~ seq, ncol = 4, scales = "free")
```

```
#ggsave(filename = "all_ASV_distributions.eps", device = "eps", dpi = 400, width = 13, height = 10, limitsize = FALSE)
```

#### Plotting bulk DNA density distributions

```
sample_data(ps_filt_RA)$name <- paste("Day", sample_data(ps_filt_RA)$Day, "-", sample_data(ps_filt_RA)$Substrate)
sample_data(ps_filt_RA)$name <- factor(sample_data(ps_filt_RA)$name, levels=rev(unique(table_for_plot.g2$name)))

ggplot(sample_data(ps_filt_RA), aes(x=Buoyant_density, y=as.numeric(paste(Fraction.DNA.proportion)), fill = name)) +
  geom_area(aes(fill=name), color = "black", position = "identity", alpha = 0.8) +
  scale_x_continuous(limits = c(1.680,1.7455), breaks = seq(1.680,1.7455,0.005)) +
  theme_classic() +
  theme(axis.text.x=element_text(angle=90, vjust=0.4, hjust=1), legend.title=element_blank()) +
  labs(x="Fraction Density (g/ml)", y="DNA in Fraction / Total") +
  guides(fill = guide_legend(reverse = TRUE)) +
  scale_fill_manual(values = c("firebrick4","firebrick2","darkorange1","lightskyblue"))
```

#### qSIP calculations using HTSSIP packahe

##### Following vignette --> https://rdrr.io/cran/HTSSIP/f/vignettes/qSIP.Rmd

```
#splitting phyloseq object into pairs of control and heavy for each of the 3 heavy timepounts
ps_D1 = prune_samples((m$Substrate=='Light' & m$Day==1) | (m$Substrate=='Heavy' & m$Day==1), ps_filt_all)
ps_D3 = prune_samples((m$Substrate=='Light' & m$Day==1) | (m$Substrate=='Heavy' & m$Day==3), ps_filt_all)
ps_D4 = prune_samples((m$Substrate=='Light' & m$Day==1) | (m$Substrate=='Heavy' & m$Day==4), ps_filt_all)

#reading in DNA quantification data from Qubit — note that I use this instead of qPCR data as in the vignette, but to use the package functions, I need to use misleading names ike "qPCR_tech_rep_mean"
quant_D1<-(read.table("DNA_quant_tab_D1.txt", header=TRUE, colClasses = c("character","character","character","character","numeric","numeric","numeric"), check.names = F, na.strings="", sep="\t"))
quant_D1$qPCR_tech_rep_mean <- quant_D1$DNA_quant * 1000 # I multiply by 1000 here because the DNA quantification values are orders of magnitude smaller than qPCR values, and to use the package I need to them in the same ballpark
quant_D3<-(read.table("DNA_quant_tab_D3.txt", header=TRUE, colClasses = c("character","character","character","character","numeric","numeric","numeric"), check.names = F, na.strings="", sep="\t"))
quant_D3$qPCR_tech_rep_mean <- quant_D3$DNA_quant * 1000
quant_D4<-(read.table("DNA_quant_tab_D4.txt", header=TRUE, colClasses = c("character","character","character","character","numeric","numeric","numeric"), check.names = F, na.strings="", sep="\t"))
quant_D4$qPCR_tech_rep_mean <- quant_D4$DNA_quant * 1000

# transforming ASV counts to relative abundance and multiplying by fraction DNA quantification values
qsip_D1 <- OTU_qPCR_trans(ps_D1, quant_D1)
qsip_D3 <- OTU_qPCR_trans(ps_D3, quant_D3)
qsip_D4 <- OTU_qPCR_trans(ps_D4, quant_D4)


#setting the replicate value to 1 for atom fraction excess calculations
sample_data(qsip_D1)$Replicate <- 1
sample_data(qsip_D3)$Replicate <- 1
sample_data(qsip_D4)$Replicate <- 1

#I modified the qSIP_atom_excess function to include the parameters for 15N, which were not inlcuded in the pacakge
source("qSIP_atom_excess_sourcecode.R")

#running atom fraction excess calculations
atomX_D1 = qSIP_atom_excess(qsip_D1,
                         control_expr='Substrate=="Light"',
                         treatment_rep='Replicate',
                         isotope = "15N")
atomX_D3 = qSIP_atom_excess(qsip_D3,
                         control_expr='Substrate=="Light"',
                         treatment_rep='Replicate',
                         isotope = "15N")
atomX_D4 = qSIP_atom_excess(qsip_D4,
                         control_expr='Substrate=="Light"',
                         treatment_rep='Replicate',
                         isotope = "15N")
df_atomX_boot_D1 = qSIP_bootstrap(atomX_D1, n_boot=100)
df_atomX_boot_D3 = qSIP_bootstrap(atomX_D3, n_boot=100)
df_atomX_boot_D4 = qSIP_bootstrap(atomX_D4, n_boot=100)

#reording tables by atom fraction excess
CI_threshold = 0
df_atomX_boot_D1 = df_atomX_boot_D1 %>%
  mutate(Incorporator = A_CI_low > CI_threshold, Day = 1,
                OTU = reorder(OTU, -A))
df_atomX_boot_D3 = df_atomX_boot_D3 %>%
  mutate(Incorporator = A_CI_low > CI_threshold, Day = 3)
df_atomX_boot_D4 = df_atomX_boot_D4 %>%
  mutate(Incorporator = A_CI_low > CI_threshold, Day = 4)

#combining tables
df_atomX_boot_all <- rbind(df_atomX_boot_D1, df_atomX_boot_D3, df_atomX_boot_D4)
df_atomX_boot_all$Day <- as.character(df_atomX_boot_all$Day)

#determining our empirical threshold for incorporation
thresh <- df_atomX_boot_all$A[df_atomX_boot_all$A < 0] %>% mean * -3

#plotting
ggplot(df_atomX_boot_all, aes(x = A, y = OTU, fill=Day)) +
  #geom_col(stat='identity', width = 0.7, position=position_dodge(0.7), color = "black") +
  geom_bar(stat='identity', width = 0.7, position="dodge", color = "black") +
  scale_fill_grey() +
  scale_y_discrete(limits = rev(levels(df_atomX_boot_all$OTU))) +
  #scale_fill_manual(values = c("darkorange1", "firebrick2", "firebrick4")) +
  #scale_fill_manual(values = c("lightskyblue","darkorange1", "firebrick2", "firebrick4")) +
  geom_hline(yintercept=0) +
  labs(x='Atom fraction excess', y='ASV') +
  theme_classic() +
  #geom_hline(yintercept=thresh, linetype="dashed") +
  geom_vline(xintercept=thresh, linetype="dashed") +
  theme(axis.title.x = element_text(vjust=-0.5), text = element_text(size=20), axis.text.x=element_text(angle=90, vjust=0.4, hjust=1, size=12))
```

```
#ggsave(filename = "Figure3_atom_fraction_excess.eps", device = "eps", dpi = 400, width = 10, height = 8, units = "in")
```
